# Blown-Arc plasma: A safe treatment with potential to boost wheat seed germination without phytotoxicity

**DOI:** 10.64898/2025.12.18.695088

**Authors:** M. Kaur, D. Hüberli, K. L. Bayliss

**Affiliations:** School of Agricultural Sciences, College of Environmental and Life Sciences, and Food Futures Institute, Murdoch University, 90 South Street, Murdoch, Western Australia 6150; Department of Primary Industries and Regional Development, 1 Nash Street, Perth, Western Australia 6000

**Keywords:** Gliding arc, cold plasma, wheat, seed treatment, atmospheric air, germination, field emergence, phytotoxicity

## Abstract

**Context:** Cold plasma is an ionised gas composed of reactive nitrogen and oxygen species, with demonstrated potential for improving wheat seed vigour.

**Aim:** To assess wheat seed vigour and grain quality after Blown-Arc plasma treatment.

**Methods:** Wheat seeds were treated with Blown-Arc plasma for 60 or 180 s from a distance of 21 cm in a closed environment to assess *in vitro* germination, seed quality, seedling emergence, and crop yield in the field.

**Key Results:** Seeds treated with Blown-Arc plasma showed significantly faster initial *in vitro* germination (day 4), although total germination by day 8 was similar to untreated controls. In the field, the treated seed showed no significant difference in seedling emergence in 2020, but in 2021 seed treated for 180 s produced a significantly lower number of seedlings, possibly due to differences in the soil microbial environment. The wheat grown from 180 s Blown-Arc plasma-treated seed produced more heads per plant; however, overall yield was unchanged. The treatment had no adverse effect on grain quality parameters, all remaining within acceptable Australian standards.

**Conclusions:** Blown-Arc plasma did not alter seedling emergence, yield, or grain quality properties of the crop produced from treated wheat seeds.

**Implications:** Blown-arc plasma is a safe treatment that can potentially increase wheat seed germination without affecting grain quality or yield. Faster germination of seeds when sown under adequate moisture may promote healthy root systems and plant growth. Further research is needed to determine how these benefits translate under variable field conditions in Australian wheat-growing regions.

## Introduction

Wheat is a staple cereal crop consumed worldwide, and with the increasing population, the demand for cereal grain is rising exponentially (Alexandratos and Bruinsma 2012; OECD-FAO 2025). As the demand grows, there is increasing interest in safe, sustainable, and chemical-free agricultural practices to maintain crop yields and meet quality standards (Chowdhuri and Pal 2025). One of the critical factors influencing crop establishment and productivity is the seed’s ability for rapid emergence and seedling establishment (Finch-Savage and Bassel 2015). At the postharvest stage, quality parameters including protein content, physical grain traits and yield, directly influence end-use suitability and grain market value (Riaz *et al*. 2024). Conventional farming depends on irrigation, fertilisation, fungicides and pesticides to improve crop yield, which impact the cost of production and leave chemical residues on the harvested seeds (De Ponti *et al*. 2012; Carvalho 2017). Therefore, environmentally safe methods that improve seed performance and crop production without compromising grain quality are being extensively studied globally. Cold plasma is an ionised gas containing reactive oxygen and reactive nitrogen species, free electrons and radicals (Whitehead 2016). Several cold plasma systems have been used to treat different plant seeds and grain, including dielectric barrier discharge, plasma jets, and gliding arc to improve seed performance (Adhikari *et al*. 2020). The effectiveness of these systems can vary depending on the plasma source, exposure time, gas composition, and the biological material being treated (Kaur *et al*. 2020). Despite the potential of cold plasma, the practical application of the treatment to improve seed performance remains inconsistent and requires additional research, particularly under field conditions.

Several studies have reported that cold plasma increases wheat seed *in vitro* germination (Šerá *et al*. 2010; Los *et al*. 2018; Roy *et al*. 2018; Los *et al*. 2019; Lotfy *et al*. 2019; Velichko *et al*. 2019). Treatment with cold plasma increases the seed’s water permeability by etching the seed coat, resulting in faster imbibition and more uniform germination (Chen *et al*. 2012; Chen *et al*. 2016). However, the impact on wheat seed vigour especially in field conditions, is unexplored. Results are often inconsistent due to differences in treatment protocols and plasma systems. For instance, Jiang *et al*. (2014) reported an increased *in vitro* germination rate, and yield in field experiments after 15 s of cold plasma treatment of seeds, but the statistical significance of the yield result was unclear. Roy *et al*. (2018) also observed an increase in germination rate *in vitro* and yield in the field, following 3 and 6 min treatments of wheat seeds, respectively. Saberi *et al*. (2022) reported a significant increase in yield, seed protein and starch content under field conditions after 180 s of treatment. Although these international studies have shown improved germination after cold plasma treatment under laboratory conditions, none, to our knowledge, have evaluated whether these effects translate to seedling emergence and grain quality under Australian field conditions.

Among cold plasma systems, Blown-Arc plasma (Enercon, Wisconsin, US) produces a gliding arc plasma using ambient air as the working gas and generates a stable plasma arc that can be applied at atmospheric pressure (Kaur *et al*. 2024). This makes it a clean, sustainable and scalable option for seed treatment. To establish baseline effects of Blown-Arc plasma on wheat seeds, this study initially investigated germination and nutrient quality of wheat seeds *in vitro*. To determine whether these effects translated to natural environments, seedling emergence, crop yield, and postharvest seed quality were further assessed under irrigated field conditions across two growing seasons. The hypothesis tested in the study was that Blown-Arc Plasma treatment has no phytotoxic effect on treated seed vigour or nutrient quality of grain produced

## Materials and Methods

### Plasma apparatus and seed treatment

Visibly clean Australian hard wheat seeds of *Triticum aestivum* cv. Scepter with a moisture content of 13% was provided by the Department of Primary Industries and Regional Development (DPIRD) Western Australia (WA). To prepare plasma treated seeds, a Blown-Arc Pro Series plasma surface treater (Enercon, Wisconsin, US) was used as described previously (Kaur *et al*. 2022). Seeds were distributed in a single layer in sterile 150 mm Petri plates, lid removed, and treated with Blown-Arc plasma for 0, 60 or 180 s at a distance of 21 cm in a closed environment (33 L plastic box), under conditions derived from the optimised plasma treatment protocol of Kaur *et al*. (2024).

### In vitro *germination assessment*

Germination tests were performed according to the International Seed Testing Association (ISTA) germination test protocol (ISTA 2020). A sample of 400 seeds was treated with Blown-Arc plasma as described above, then 50 seeds from each treatment sample were aseptically transferred into germination plates, 150 mm Petri plates with Whatman filter paper soaked with 2 mL of sterile distilled water (pH = 6.9). The plates were sealed with parafilm and incubated at 20 °C in the dark for eight days. Seeds were examined after four and eight days of incubation. The seeds were considered germinated when the length of the coleoptile and radicle reached 1 mm. After the germination count on day four, all the germinated seeds were removed from the plates, and 1 ml of sterile distilled water was added to each plate to maintain the moisture levels (ISTA 2020). Data from days four and eight were combined to give the total germination, and the germination rate was assessed from the proportion of total germination that occurred by day four. The experiment was conducted twice.

### Wheat seed quality assessment

For nutrient quality analysis, a larger sample size comprising 150 g of seeds was distributed uniformly in a single layer in a 28 x 20.5 cm plastic tray and treated with Blown-Arc plasma for 60 and 180 s in the closed environment, as described above. No plasma treatments were applied to the controls; each treatment and control consisted of ten replicate samples. After treatment, all samples were immediately aseptically transferred into Ziplock bags. The samples were delivered to the Australian Export Grain Innovation Centre (AEGIC, 3 Baron-Hay Ct, South Perth, WA 6151), where they were stored at 15 °C and 35% relative humidity. The quality assessments were conducted within seven days of storage. Near-infrared reflectance spectroscopy was performed using FOSS XDS NIR systems (Hillerod, Denmark) to estimate flour yield, particle size and water absorption, indicating seed hardness and colour index on the scale of Minolta b* to measure yellowness of the seeds and wheat proteins (Blakeney *et al*. 2011). The falling number test was performed to measure the samples’ alpha-amylase enzyme activity or starch strength using a Perten Falling Number 1700 System (PerkinElmer, Waltham, MA). The quality assessment experiment was repeated.

### Field trials and conditions

The field trials in 2020 and 2021 were conducted in two irrigated screen houses of 320 m^2^ at DPIRD WA (3 Baron-Hay Ct, South Perth, WA 6151). The screen houses were covered with fine mesh cloth that helped retain moisture and heat while allowing natural airflow, light exposure, and rainwater to pass through. Before establishing the trials, soil was collected from a depth of 0-10 and 10-30 cm from the front, middle and back sections of the screen house and analysed (CSBP Soil and Plant Analysis Laboratory, Bibra Lake, WA 6163) for soil texture, pH, salinity and total nutrients (Table S1). The soil analysis showed that although the available nitrogen in the screen house soil used in 2021 was higher than in 2020, it was significantly lower than the adequate levels of 10-50 mg/kg in both years (Table S1). Every four to six weeks, a nitrogen-phosphorus-potassium blend (Crop builder N9, Nutrien Ag Solutions, Midvale, Western Australia) was applied to all trials at a rate of 100 kg/ha in both years; thus, the nitrogen nutrition was adequate in both years.

To assess the performance of treated seeds, two samples of 400 seeds were treated with Blown-Arc plasma as described above and were transferred into plastic Ziplock bags allocated for that treatment, mixed and sown on the same day. A control seed treatment with a fluquinconazole fungicide, Jockey Stayer (Bayer, Australia), was also used to compare the results of Blown-Arc plasma-treated seeds with a fungicide treatment commonly used in Australia to control seed-borne diseases of wheat (e.g., common bunt and loose smut). For the fungicide treatment, 250 g seeds were treated using a 36.5 cm rotary seed coater (BraceWorks, Canada) at an application rate of 4.5 L/T seeds diluted with 1.5 L water. Two samples of 400 fungicide treated seeds were stored in Ziplock plastic bags at 4 °C for two weeks before sowing the 2020 trials. Another batch of two samples of 400 fungicide-treated seeds was stored at 4 °C for eleven months to be used for 2021 trials. The seeds were mixed and brought to room temperature 24 h before sowing. Fungicide-treated seeds were sown on the same day as those treated with Blown-Arc plasma.

There were 32 plots in each field trial, consisting of four treatments: the untreated control, 60 or 180 s cold plasma treatment, and fungicide treatment, and eight replicate plots. The seeds were hand-sown in late June 2020 and mid-May 2021, in 1 x 1 m plots with 100 seeds per plot, with rows 10 cm apart. A 1 m wide buffer between the plots was sown with a row of oats (cv. Williams) in the middle of all buffers. Plots for both trials were arranged in a randomised block design (Fig. S1).

All plots in the 2020 trials were irrigated using a centralised irrigation system at ~8 mm/h. Plots were watered twice weekly for 1 h in the morning when there was little or no rainfall (mid-September 2020 onwards). The screen house used for the trials in 2021 had overhead sprinklers delivering 0.63 – 0.76 L/min, and the plots were irrigated for 4 h per day. In 2021, plants exhibited mildew infection at Zadoks Growth Stage 53-57 (Poole 2005); this was controlled using 3 mL/L of a pyrimidine-based fungicide, Nimrod (Adama Ltd, St Leonards, New South Wales).

### Seedling emergence and harvest

The number of seedlings that emerged was recorded in each plot fourteen days after sowing. Two weeks before harvest, the number of wheat heads per plant in 20 randomly selected plants from the yield trial plots were recorded.

Wheat heads were hand harvested from both trials when the seeds were hard to crush between fingers (Zadoks Growth Stage 92 – 93) and collected in separate paper bags for each plot and stored at 37 °C for two weeks to remove any moisture before threshing. In 2020, threshing was completed within four weeks after dry incubation using a mechanical thresher (Simpson Pope Ltd, Beverley, Adelaide), whereas a relatively newer mechanical thresher (Wintersteiger Inc., Salt Lake City, US) was used for the 2021 harvest, which was completed within a week after harvest. After threshing, seeds were stored at 4 °C for a week in both years then analysed for total yield quality properties.

### Postharvest yield assessments

The total yield for both years in the yield trial was determined by weighing the harvested seeds from each plot. The 1000 seed weight was measured using a seed counter (Contador, Pfeuffer GmbH, Germany). The percentage of unmillable seeds was determined from a 0.5 L seed sample using a chondrometer (Graintec Scientific, Toowoomba, Queensland). The sample was processed through a 2.0 mm sieve on an agitator (Graintec Scientific, Toowoomba, Queensland) and the weight of seeds falling through the sieve was recorded. The percentage of unmillable screenings was calculated by putting the weight of seeds that fell through the sieve over the total test weight that went into the sieve. The moisture content was also measured using the Grist Marconi moisture meter TF 933C (Marconi Instruments Ltd, UK) by the AEGIC.

### Statistical analysis

Data were expressed as means and standard errors of all replicate plots with the same treatment and were analysed using R software (version 4.0.2) (R Core Team 2013). Before the analysis, the mean of all the samples was calculated from all replicates in a treatment. Differences between the Blown-Arc plasma treatments were determined using ANOVA. Mean differences between treated and untreated samples were compared by Fischer’s LSD in R. For the data from *in vitro* experiments, the variance between the repeated trials was checked using Levene’s test and was not significantly different; therefore, the data from repeated trials were combined before further mean comparison analysis. Square root data transformation was performed on all data showing heterogeneity of variance, except nutrient quality data, which was analysed without transformation.

For the field trials, Levene’s test of Variance between the trials in 2020 and 2021 indicated a significant difference between years (P < 0.001) for all data except for yield and quality, which could be combined. Therefore, the data for seed emergence and the number of wheat heads per plant were analysed separately each year. The difference in the treatments was considered statistically significant at P < 0.05. The results were plotted using the ggplot2 package (Wickham and Sievert 2016).

## Results

### In vitro *germination and grain quality*

The initial germination rate (day four) of seeds treated with Blown-Arc plasma for 60 and 180 s increased significantly (P = 0.01) compared with untreated controls. However, the total germination percentage by day eight did not differ from the controls (Table 1).

**Table 1.**
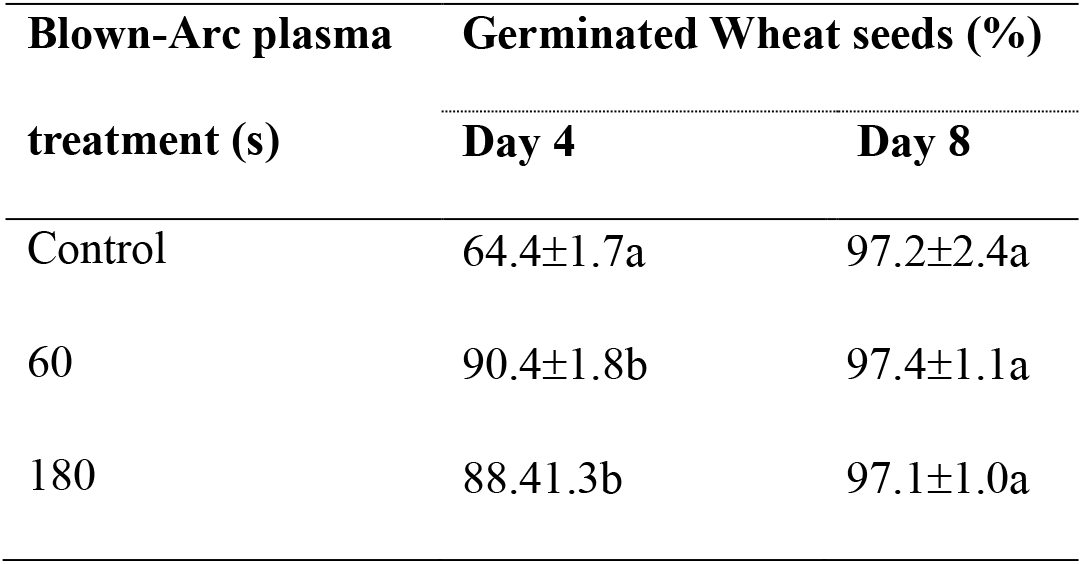
Germination of wheat seeds, untreated control and treated with Blown-Arc plasma for 60 and 180 s. The mean percentages and standard errors of germinated wheat seeds on day eight, and the proportion germinated by day four (expressed as a percentage of the total germination) are shown. Means for the percent germination within a column with the same letter are not significantly different (P < 0.05, Fisher’s LSD).

Blown-Arc plasma treatment had no impact on the flour yield or seed quality as assessed by the falling numbers of the seeds. Grain particle size of treated seeds was significantly (P < 0.001) increased, and water absorption decreased significantly after the Blown-Arc plasma treatment for 180 s (Table 2). However, these parameters remained within the acceptable values of Australian grain trade standards (GTA 2022).

**Table 2.** Means and standard errors of seed quality parameters-falling numbers, flour yield, particle size index, water absorption, colour index and wheat protein following the Blown-Arc plasma treatments of wheat seeds for 60 and 180 s, and untreated control. Means are of 20 samples, parameters with the same letter are not significant (P < 0.05, Fisher’s LSD).

| Blown-Arc<br>plasma<br>treatment<br>(s) | Falling<br>numbers (s) | Flour yield<br>(%) | Particle size<br>index (PSI) | Water<br>absorption<br>(%) | Colour<br>index<br>(Minolta<br>b*) | Wheat<br>protein (%) |
| --- | --- | --- | --- | --- | --- | --- |
| Control | 439.1±4.1a | 72.0±0.2a | 11.8±0.3c | 61.2±0.3a | 11.1±0.1a | 11.0±0.1a |
| 60 | 446.9±4.7a | 71.8±0.2a | 13.2±0.3b | 60.7±0.3a | 11.2±0.1a | 10.9±0.1a |
| 180 | 446.3±11.0a | 72.3±0.2a | 16.0±0.3a | 60.0±0.3b | 11.3±0.1a | 10.9±0.05a |

### Seedling emergence in the field, yield and grain quality

In 2020, seeds treated with Blown-Arc plasma or fungicide did not show any difference in seedling emergence fourteen days after sowing. However, in 2021, 180 s Blown-Arc plasma treated seed had significantly reduced (P = 0.05) seedling emergence, while the fungicide treatment increased seedling emergence (Fig. 1). In both years, plants grown from 180 s Blown-Arc plasma treated seeds had a higher number of heads per plant than the control, but the increase was statistically significant (P = 0.05) in 2020 only. In 2021, the fungicide-treated seed produced a significantly lower (P < 0.05) number of heads (Fig. 2).

**Fig. 1.**
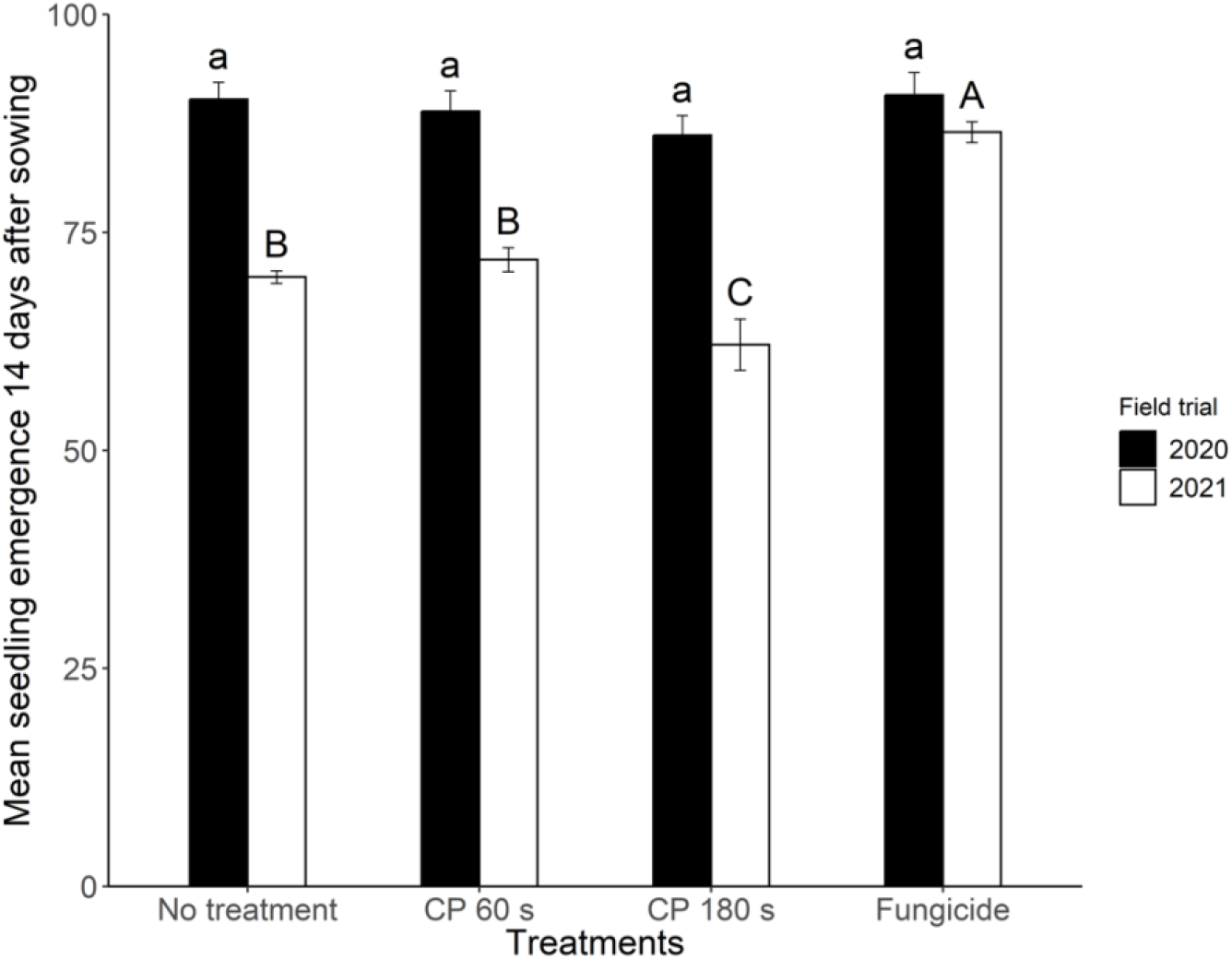
The number of emerged seedlings fourteen days after sowing seeds in four pre-sowing treatments: no treatment controls, Blown-Arc plasma (CP) treatment for 60 or 180 s, and fungicide treatment. There were 100 seeds per treatment. Bars indicate standard errors of the means (n = 8). Lower-case letters represent the mean of 2020 samples, and upper-case letter shows the mean of 2021 samples. Means with the same letter are not significantly different across all treatments within a year (P = 0.05, Fisher’s LSD).

**Fig. 2.**
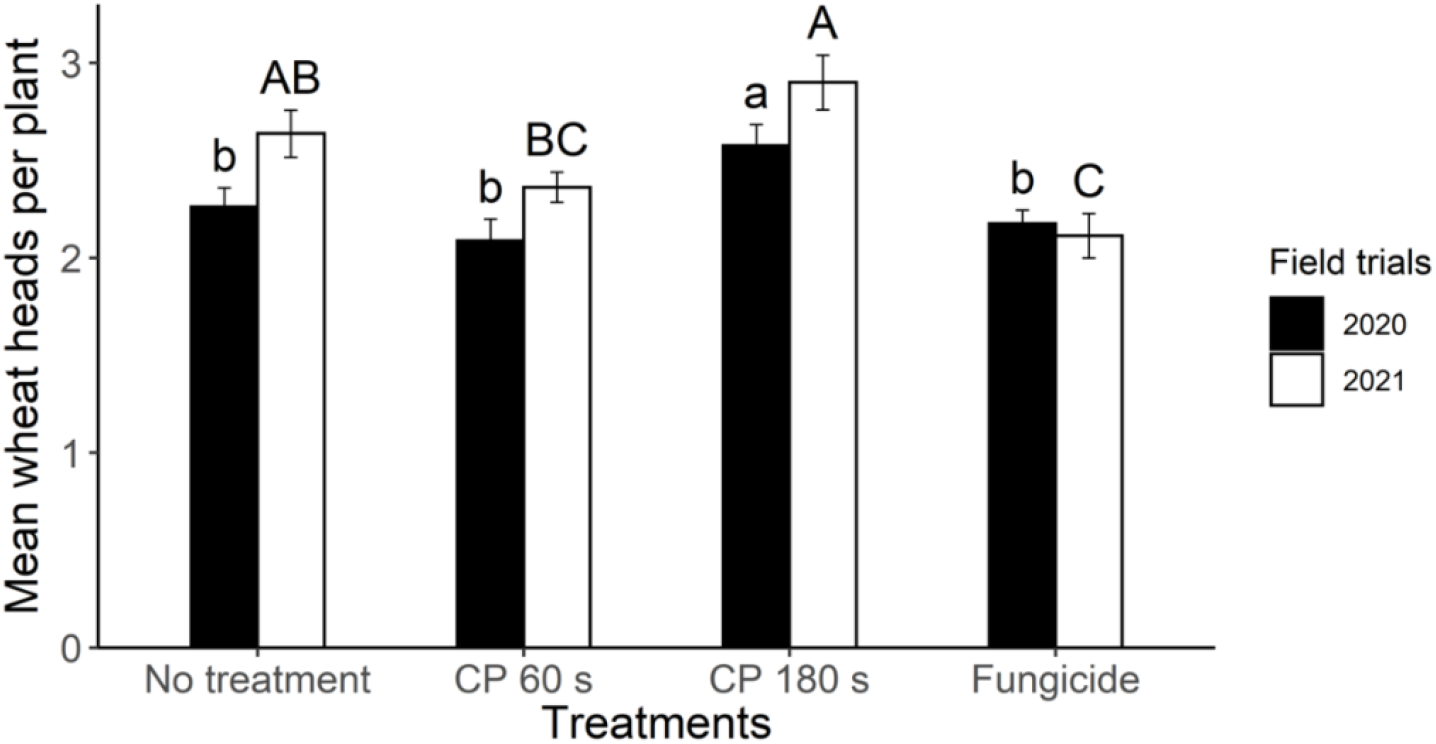
The mean number of wheat heads per plant recorded from yield trials in 2020 and 2021. Before sowing, seeds were treated with 60 or 80 s of Blown-Arc plasma (CP) or a fungicide. Bars indicate standard errors of the means (n = 8). Different lower-case letters indicate a significant difference between treatments (P = 0.05, Fisher’s LSD) in 2020, and upper-case letters show differences between treatments in 2021.

The total yield, unmillable screenings, 1000 seed weight, and moisture content of seeds from the yield trials showed no significant differences between controls and plots sown with Blown-Arc plasma or fungicide treated seeds in either year (Table 3).

**Table 3.** Means of grain quality properties harvested from grain treated with Blown-Arc plasma (CP) for 60 or 180 s or with fungicide from the yield trials in 2020 and 2021. Means for each parameter with the same letter are not significant (P = 0.05, Fisher’s LSD).

| Grain treatment | Total yield<br>(t/ha) | Unmillable<br>screenings (%) | 1000 seed<br>weight (g) | Moisture<br>content (%) |
| --- | --- | --- | --- | --- |
| No treatment<br>control | 4.0 a | 0.7 a | 55.2 a | 11.1 a |
| 60 s CP | 4.0 a | 0.6 a | 56.1 a | 11.0 a |
| 180 s CP | 3.9 a | 0.6 a | 55.6 a | 11.0 a |
| Fungicide | 4.1 a | 0.5 a | 56.1 a | 11.0 a |

## Discussion

This study found that the Blown-Arc Plasma is a safe treatment for wheat seed and does not change seed vigour *in vitro* and in field. Longer plasma treatment increased early *in vitro* germination up to day four, suggesting a stimulatory effect on early seed metabolism, consistent with other *in vitro* studies. For example, the germination rate increased after 15 s (Velichko *et al*. 2019) and 30 s ((Los *et al*. 2018) treatments, but 45 and 180 s were detrimental ((Los *et al*. 2018; Velichko *et al*. 2019). Plasma treatments are known to improve water uptake, possibly increasing metabolic activation and enzyme activity required for germination (Roy *et al*. 2018; Los *et al*. 2019; Lotfy *et al*. 2019). The reactive species generated by cold plasma promote the growth of roots and shoots, leading to a faster onset of germination (Iranbakhsh *et al*. 2017). Similar observations to our study were reported by Šerá *et al*. (2010), stating higher germination in 3 min plasma-treated wheat seeds on day four, with no difference by day eight or twelve. They observed no difference in germination for longer plasma treatment. The current study supports the general trend with *in vitro* studies that short to moderate plasma treatments may stimulate early germination in wheat, but extended treatment, such as the 180 s in this study, does not improve final germination. Together, these results show the importance of treatment duration for cold plasma, and the need for careful optimisation of plasma parameters due to the complex nature of plasma systems.

The influence of Blown-Arc plasma treatment on seed performance and yield were compared to the standard Australian practice of utilising fungicide-treated seeds (DPIRD 2024). Fungicide-treated seeds showed consistent crop establishment in both years, suggesting effective and uniform protection. However, varied seedling emergence in the control and plasma treated seed in both years could be linked to differences in the soil microbial environment. Microbiome analysis of wheat harvested from the same field for both years showed differences in fungal abundance and diversity, likely reflecting changes in the fungal communities (Kaur *et al*. 2024). Despite variation in seedling emergence among the treatments, the overall yield remained unchanged. Interestingly, Blown-Arc plasma treated seeds produced a higher number of wheat heads per plant despite fewer emerged seedlings, possibly explaining why the yield was not different. This finding may seem counterintuitive at first glance, as one might expect that having more plants would lead to a greater yield. However, plants often compensate for lower crop density by producing more productive tillers (Kiss *et al*. 2018). It is also possible that the increased number of heads was influenced by the stimulatory effects of the cold plasma treatment; however, further comparative research with different cultivars and broader seed treatments, including biological and chemical, is required to evaluate its stimulatory impact on seed performance and yield.

Previous studies have suggested that cold plasma treatment can positively impact wheat yield under field conditions, but this was not observed in the current study. Jiang *et al*. (2014) observed a 5.89% increase in yield after a 15 s treatment using a customised helium-based plasma apparatus, while Roy *et al*. (2018) found a significant yield increase of about 20% following a 6 min treatment with dielectric barrier discharge plasma systems. Saberi *et al*. (2022) reported a significant increase of nearly 20% in yield, and ~10% in protein and starch content after a 180 s treatment with a radio frequency plasma system. Filatova *et al*. (2020) also noted a 2.3% increase in yield after 5 min treatment using a customised capacitively coupled plasma system, though it is unclear if the increase was significant. These studies used plasma systems and treatment conditions that differ substantially from the Blown-Arc plasma system used in the current study, which may explain the variations.

Australian hard wheat has an excellent reputation on the international market for good flour colour, high flour yield and water absorption properties, and is highly suited for pan breads, flat breads, yellow alkaline noodles, and steamed breads (Blakeney *et al*. 2011). Chaple *et al*. (2020) suggested that cold plasma treatment could influence some of the functional properties of the flour obtained from treated wheat seeds, but the treatments could be optimised for their beneficial effect. Their study showed that protein structure and water absorption properties of wheat seeds remained unchanged, suggesting the essential quality parameters were maintained after the treatment. In the current study, the grain was measured according to Australian quality standards, and despite minor alterations in the seed hardness and colour index observed after Blown-Arc plasma treatment, the values remained within the acceptable standards (GTA 2022). Thus, treatment may be used without compromising the high-quality reputation of Australian grain.

## Conclusion

This study found that Blown-Arc plasma treatment accelerated the *in vitro* germination of wheat seeds but this did not translate to improved seedling emergence in the field. Blown-Arc plasma treatment did not change the seed quality parameters of flour yield, wheat protein or falling numbers of the treated seeds. The plants grown from the seeds with the 180 s plasma treatment had an increased number of heads per plant, but overall yield remained unchanged. This study demonstrated that cold plasma does not have a phytotoxic effect on the treated wheat seed. More research is needed to understand how the observed increase in wheat heads per plant might translate into increased yield, focusing on evaluating cold plasma technologies under variable field conditions in Australian wheat-growing regions.

## Data availability

The data supporting this study will be shared upon request.

## Acknowledgments

The Department of Primary Industries and Regional Development WA supported the first author for providing financial support under the Boosting Grain, Research and Development Postgraduate Scholarships program. The first author is thankful to Dr Manjree Agarwal and Dr Bill Dunstan for providing technical advice on the moisture content adjustment of wheat seeds. Thanks to Mr Jason Bradley from DPIRD for technical support with the field trials. Thanks to Dr Wendy Hunt from the Australian Export Grain Innovation Centre for her assistance with grain quality analysis and Em. Prof. Jen McComb for her valuable feedback, which helped improve the study’s context.

