## Supplementary for "Blown-Arc plasma: A safe treatment with potential to boost wheat seed germination without phytotoxicity"

**Table S1** Analysis of the soil in the shade houses used in the trials of 2020 and 2021. Data from CSBP Soil and Plant Analysis Laboratory Bibra

| **Trial** | **Field location of collected samples** | **Depth** | **Colour** | **Gravel (%)** | **Texture** | **Ammonium Nitrogen (mg/kg)** | **Nitrate Nitrogen (mg/kg)** | **Phosphorus Colwell (mg/kg)** | **Potassium Colwell (mg/kg)** | **Sulfur (mg/kg)** | **Organic Carbon (%)** | **Conductivity (dS/m)** | **pH Level (CaCl2)** | **pH Level (H2O)** |
| --- | --- | --- | --- | --- | --- | --- | --- | --- | --- | --- | --- | --- | --- | --- |
| 2020 | Front | 0-10 | BRBK (Brown Black) | 0 | 1.0 (Sand) | 1 | 1 | 67 | 38 | 0.9 | 0.31 | 0.024 | 6.2 | 7.2 |
|  | Middle | 0-10 | BRBK | 0 | 1.0 | < 1 | 1 | 59 | 31 | 3.4 | 0.33 | 0.029 | 6.4 | 7.3 |
|  | Back | 0-10 | BRBK | 0 | 1.0 | < 1 | < 1 | 56 | 35 | 4.9 | 0.32 | 0.025 | 6.3 | 7.3 |
|  | Front | 10-30 | BRBK | 0 | 1.0 | < 1 | 1 | 64 | 33 | 1.8 | 0.31 | 0.018 | 5.6 | 6.7 |
|  | Middle | 10-30 | BRBK | 0 | 1.0 | 1 | < 1 | 60 | 31 | 1.1 | 0.30 | 0.019 | 5.9 | 6.9 |
|  | Back | 10-30 | BRBK | 0 | 1.0 | 1 | 1 | 47 | 33 | 2.0 | 0.31 | 0.024 | 6.1 | 7.1 |
| 2021 | Front | 0-10 | GRBK (Grey Black) | 0 | 1.0 | 1 | 7 | 53 | 40 | 8.9 | 0.44 | 0.034 | 5.0 | 5.9 |
|  | Middle | 0-10 | DKBR (Dark Brown) | 0 | 1.0 | 1 | 8 | 51 | 37 | 7.2 | 0.36 | 0.036 | 5.1 | 6.2 |
|  | Back | 0-10 | BRBK | 0 | 1.0 | 1 | 8 | 51 | 32 | 7.3 | 0.31 | 0.039 | 5.2 | 6.1 |
|  | Front | 10-30 | BRBK | 0 | 1.0 | < 1 | 4 | 46 | 33 | 5.4 | 0.26 | 0.029 | 5.0 | 6.1 |
|  | Middle | 10-30 | BRBK | 0 | 1.0 | < 1 | 10 | 43 | 29 | 7.1 | 0.24 | 0.037 | 4.9 | 5.9 |
|  | Back | 10-30 | BRBK | 0 | 1.0 | < 1 | 11 | 50 | 24 | 4.2 | 0.24 | 0.032 | 4.7 | 5.6 |

**Yield trial**

No grain treatment

60 s cold plasma

180 s cold plasma

Fungicide treatment

Oat buffer

Central sprinkler system 2020

Overhead rotary sprinkler 2021

Screen house boundary

Soil collection points

8 m

40 m

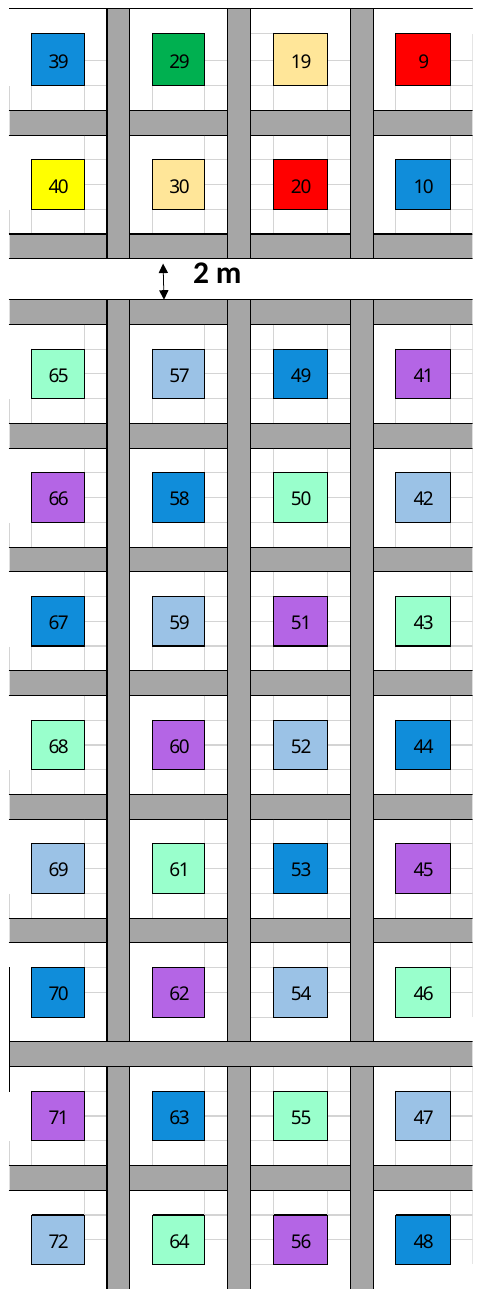

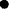

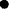

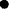

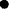

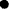

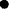

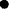

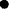

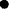

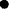

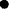

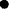

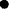

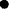

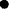

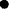

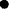

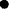

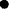

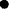

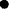

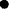

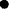

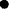

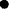

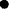

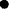

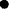

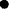

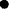

2 m Unsown buffer

Cold Plasma treated wheat trial

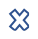

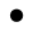

#### **Fig. S1** Field trials design for 2020 and 2021 showing randomised thirty two plots of yield trial, oats buffer zone around the plots, irrigation systems used in 2020 and 2021, location of soil collected for the soil analysis and the edges of the screen house. The number in the coloured boxed represents the plot numbers.

Front of the screen house

Space for a separate trial
